## Supplemental Figures for "Spatially displaced excitation contributes to the encoding of interrupted motion by the retinal direction-selective circuit"

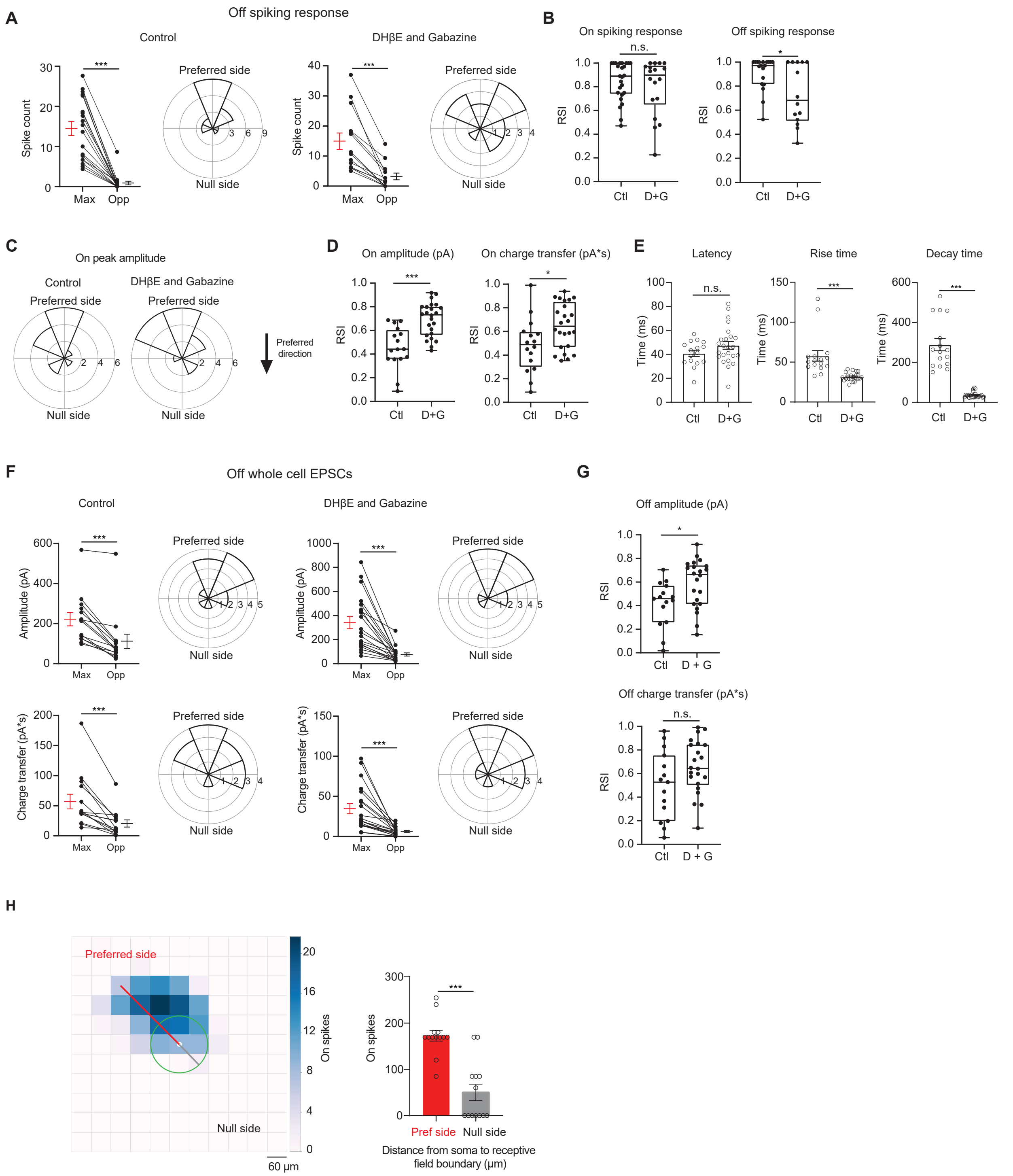


**Supplemental Figure 1. pDSGCs have spatially asymmetric RF.**

**(A)** Left: Pairwise comparison of Off mean spike counts in regions evoking the maximum number of spikes (Max) and the opposite region (Opp) in the control condition (25 cells) and polar histogram of Max region aligned to the preferred-null motion axis in the control condition. Radius indicates number of cells. Right: same as left, but in the DHβE + Gabazine condition (18 cells).**(B)** Left: Regional Selectivity Index (RSI, defined as (Max – Opp)/(Max+Opp)) of On spiking responses in the control (Ctl, 25 cells) and DHβE + Gabazine (D+G, 18 cells) conditions (p = 0.29). Right: Same as **left**, but for the Off spiking response (*p = 0.011). **(C)** Polar histograms of Max region locations determined by On EPSC peak amplitude aligned to the preferred-null motion axis in the control (left) and DHβE + Gabazine (right) conditions. Radius indicates number of cells. **(D)** RSI of On EPSC charge transfer and peak amplitude in control condition (15 cells) and DHβE + Gabazine (24 cells) conditions (charge transfer, *$p = 0.02$). **(E)** Left: Latency of On EPSC response in control condition and DHβE + Gabazine conditions ($p = 0.15$). Middle: Same as **left**, but for rise time (10% - 90%) (***$p < 0.001$). Right: Same as **left** but for decay time (90% - 10%) (***$p < 0.001$). **(F)** Top left: Pairwise comparison of Off charge transfer responses in the Max and Opp regions and the polar histograms of Max regions aligned to the preferred-null motion axis in the control condition. Radius indicates number of cells. Bottom left: Pairwise comparison of Off amplitude responses in the Max and Opp regions and the polar histograms of Max regions aligned to the preferred-null motion axis in the control condition. Top right: Same as top left, but in DHβE + Gabazine. Bottom right: Same as bottom left, but in DHβE + Gabazine. Summary statistics are mean ± SEM, ***$p < 0.001$ except where specified otherwise. **(G)** Same as **D** but for Off EPSCs (amplitude *$p = 0.014$, charge transfer $p = 0.069$). **(H)** Left: Spiking receptive field color map to 60 μm flashing spots. White circle represents soma. Green circle represents average pDSGC dendritic span. Right: Pairwise comparison of the distance to the receptive field boundary on the preferred and null sides (13 cells, ***$p < 0.001$).

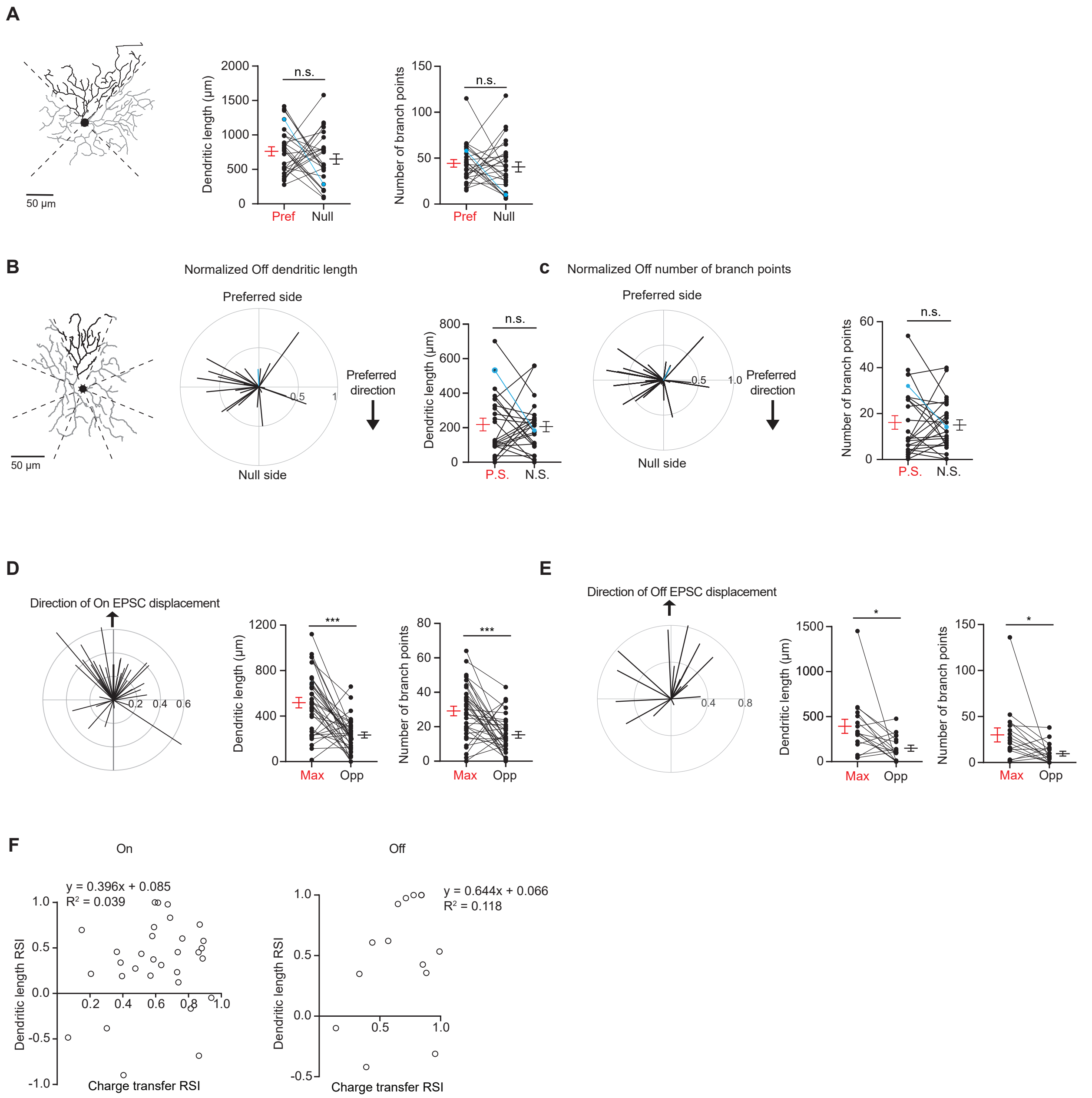


**Supplemental Figure 2. Dendritic morphology shows spatial bias along glutamatergic receptive field.**
**(A)** Left: Example morphology of a pDSGC On layer divided into quadrants. Middle: Pairwise comparison of dendritic length on the preferred vs null sides of each cell (26 cells, $p = 0.33$). Right: Total number of branch points on the preferred vs null sides of each cell (26 cells, $p = 0.25$). Example cell in blue. **(B)** Left: Example morphology of pDSGC Off layer divided into eighths. Middle**:** Normalized vector sum of Off dendritic length aligned to pDSGCs' preferred direction motion. Right: Pairwise comparison of Off dendritic length on the preferred vs null sides of each cell (24 cells, $p = 0.78$). **(C)** Same as **B** but for number of Off branch points (24 cells, $p = 0.73$).
**(D)** Left: Normalized vector sum plot of On dendritic length in eight sectors aligned to the region evoking maximal glutamatergic EPSC (upward arrow). Middle: Total dendritic length in the region evoking maximal glutamatergic EPSC as determined by peripheral spot stimulus (Max) and opposite region (Opp) (31 cells, ***$p < 0.001$). Right: Total number of branch points in the Max and Opp regions (31 cells, ***$p < 0.001$). **(E)** Same as in **D**, but for Off layer (17 cells, dendritic

length *p = 0.018, branch points *p = 0.027). **(F)** Left: On dendritic length RSI versus On EPSC charge transfer RSI (31 cells). RSI is based on regions evoking maximal glutamatergic charge transfer responses and opposite regions. Right: Same as left, but for Off dendrites and EPSCs (14 cells). Summary statistics are mean ± SEM.

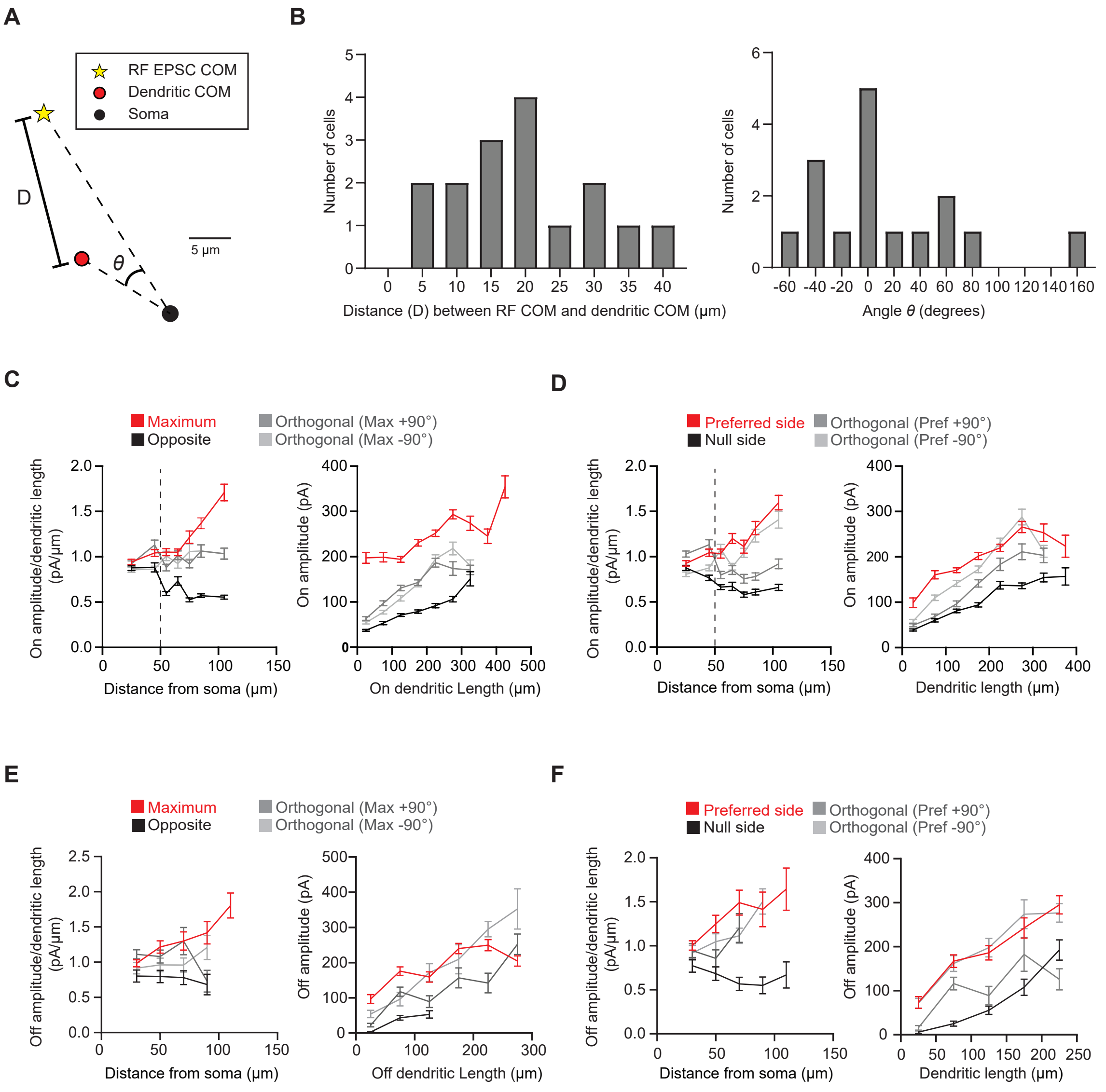


**Supplemental Figure 3. pDSGC glutamatergic synaptic excitation is displaced relative to the dendritic field.**
**(A)** The dendritic center of mass (red circle) and the RF EPSC charge transfer center of mass (yellow circle) of an example pDSGC. **(B)** Left: Histogram of the distance (D) between the dendritic center of mass and the EPSC charge transfer center of mass of each pDSGC. Right: Histogram of the angle difference (θ) between the EPSC charge transfer center of mass and the dendritic center of mass). D and θ values are illustrated in **A**. **(C)** Left: Ratio of amplitude per dendritic length versus distance from soma in the region evoking the maximum glutamatergic EPSC response (Maximum) and the opposite region (Opposite) (left, 16 cells, ***$p < 0.001$) as well as the orthogonal regions. Right: Quantification of amplitude per dendritic length in those regions for spots presented at least 50 $\mu$m away from the soma (right, 16 cells, ***$p < 0.001$). **(D)** Same as **C** but for the preferred-null motion axis. (Amplitude/dendritic length vs distance **$p = 0.001$, amplitude vs dendritic length ***$p < 0.001$). **(E)** Same as C except for Off dendrites and EPSCs (9 cells, right and left ***$p < 0.001$). **(F)** Same as D but for Off dendrites and EPSCs (9 cells, right and left ***$p < 0.001$). Summary statistics are mean ± SEM.

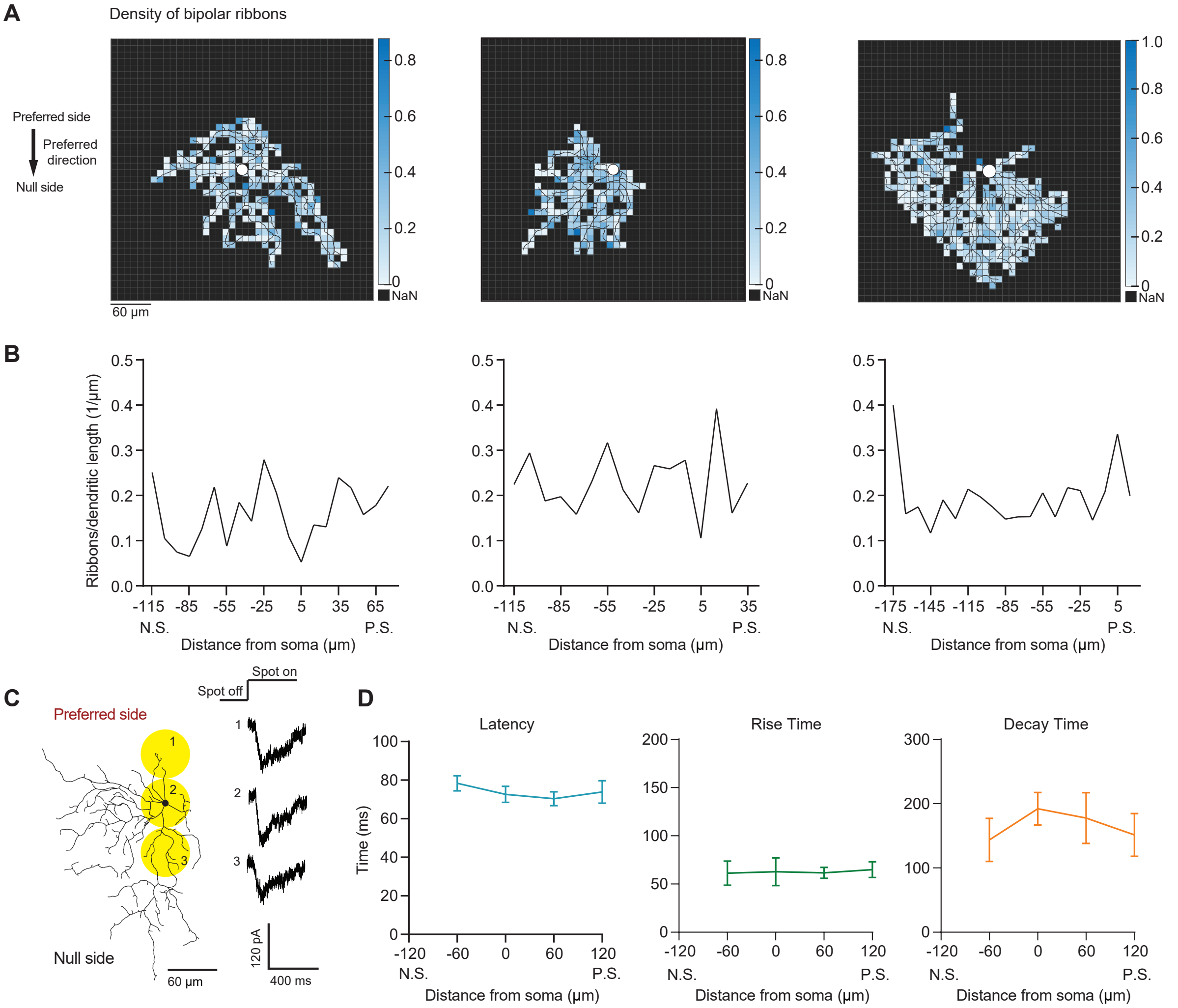


**Supplemental Figure 4. Bipolar ribbon density does not consistently change across the preferred-null motion axis.**

**(A)** Density map of bipolar ribbon synapses for 3 example cells based on the published connectomic dataset (Ding et al., 2016). Each square is 10 x 10 µm. **(B)** Quantification of ribbon density across the preferred-null motion axis. The soma location is at 0. (N.S. = null side, P.S. = preferred side). **(C)** Example glutamatergic EPSC responses to spots presented on the preferred and null sides of a pDSGC in DHβE. **(D)** Summary plots of latency ($p = 0.12$), rise time (10% - 90%; $p = 0.85$), and decay time (90% - 30%; $p = 0.93$) of glutamatergic EPSC responses along the preferred-null motion axis (N.S. = null side, P.S. = preferred side)(9 cells). Summary statistics are mean ± SEM.

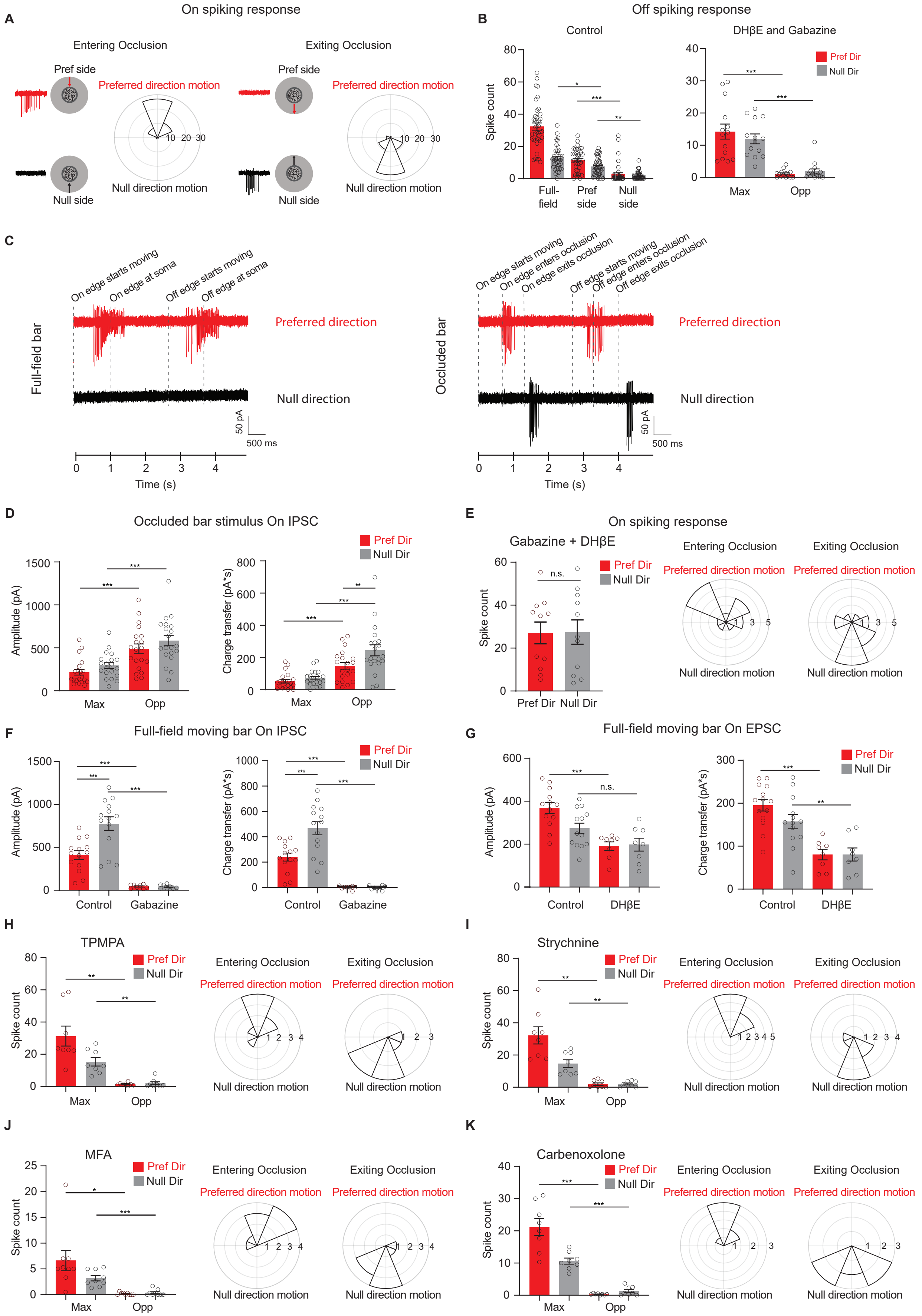
A
On spiking response
Entering Occlusion
Exiting Occlusion
Pref side
Preferred direction motion
Null side
Null direction motion
B
Off spiking response
Control
DHβE and Gabazine
Spike count
Full-field
Pref side
Null side
Max
Opp
Pref Dir
Null Dir
C
On edge starts moving
On edge at soma
Off edge starts moving
Off edge at soma
On edge enters occlusion
On edge exits occlusion
Off edge enters occlusion
Off edge exits occlusion
Full-field bar
Occluded bar
Preferred direction
Null direction
50 pA
500 ms
Time (s)
D
Occluded bar stimulus On IPSC
Amplitude (pA)
Charge transfer (pA*s)
E
On spiking response
Gabazine + DHβE
n.s.
F
Full-field moving bar On IPSC
Control
Gabazine
G
Full-field moving bar On EPSC
DHβE
H
TPMPA
I
Strychnine
J
MFA
K
Carbenoxolone

**Supplemental Figure 5. Displaced glutamatergic excitation contributes to null-direction responses in the preferred region.**

**(A)** Polar histograms of spiking vector sum locations when the bar enters the occlusion (left) versus when the bar exits the occlusion (right). Radius indicates number of cells. (48 cells). Responses are aligned to the preferred direction, which points to the top. **(B)** Left: Mean Off spike counts of pDSGCs to the full-field moving bar and the occluded moving bar stimulus on the preferred side and null side (39 cells, Full-field null dir. vs preferred side null dir. **p = 0.01, preferred side null. dir. vs null side null. dir. **p = 0.005). Right: Mean Off spike counts in DHβE + Gabazine to the occluded bar stimulus in the region evoking the maximum spiking (Max) and the opposite region (Opp) (14 cells). **(C)** Left: Example spiking traces to the full-field moving bar. Right: Example spiking traces to the occluded bar stimulus. **(D)** Mean On IPSC peak amplitude (left) and charge transfer (right) to the occluded bar in the region evoking the maximum EPSC response (Max) and the opposite region (Opp) (20 cells). **(E)** Left: Mean spike counts to the full-field moving bar in DHβE + Gabazine (11 cells, p = 0.84). Middle and Right: Polar histograms of spiking vector sum locations when the bar enters the occluder and exits the occluder. Radius indicates number of cells. (14 cells). **(F)** Mean On IPSC peak amplitude (left) and charge transfer (right) to a full-field bar in the Control (14 cells) and Gabazine conditions (8 cells). **(G)** Mean On EPSC peak amplitude (left) and charge transfer (right) to a full-field bar in the Control (13 cells) and DHβE conditions (8 cells). For amplitude: Control null dir. vs DHβE null dir. p = 0.099. For charge transfer: Control null dir. vs DHβE null dir. **p = 0.003. **(H)** Left: Mean spike counts to the bar entering and exiting the occluder in TPMPA (right, 8 cells, Pref Dir – Max vs Pref Dir – Opp **p = 0.0037. Null Dir – Max vs Null Dir – Opp **p = 0.0022). Middle and Right: Polar histograms of spiking vector sum locations when the bar enters the occluder and exits the occluder in TPMPA Radius indicates number of cells. (8 cells). **(I)** As in **H**, but in strychnine (8 cells, Pref Dir – Max vs Pref Dir – Opp **p = 0.0012. Null Dir – Max vs Null Dir – Opp **p = 0.0017). **(J)** As in **H**, but in MFA (9 cells, Pref Dir – Max vs Pref Dir – Opp *p = 0.021). **(K)** As in **H**, but in Carbenoxolone (8 cells). Summary statistics are mean ± SEM, ***p < 0.001 except where specified otherwise.

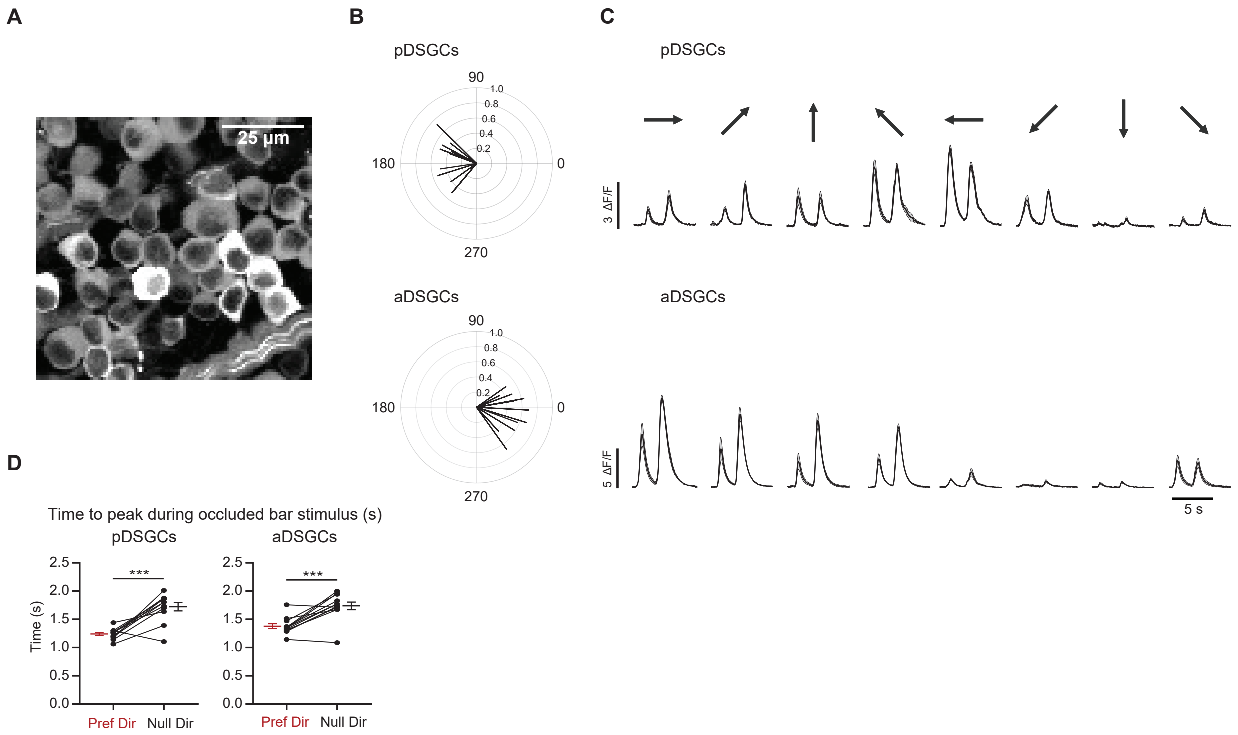


**Supplemental Figure 6. Calcium imaging of anterior and posterior-preferring On-Off DSGCs.**
**(A)** Z-stack standard deviation projection of GCaMP6f-expressing cells in the ganglion cell layer. **(B)** Top: Vector sum plot of peak calcium signal amplitudes of pDSGCs. Bottom: Vector sum plot of peak amplitudes of aDSGCs. **(C)** Example calcium traces of a pDSGC (top) and an aDSGC (bottom) to full-field moving bar in eight directions. **(D)** Left: Preferred direction time to peak vs null direction time to peak during the occluded bar stimulus for pDSGCs (12 cells, ***p < 0.001). Right: Same as top but for aDSGCs (12 cells, ***p < 0.001). Summary statistics are mean ± SEM.

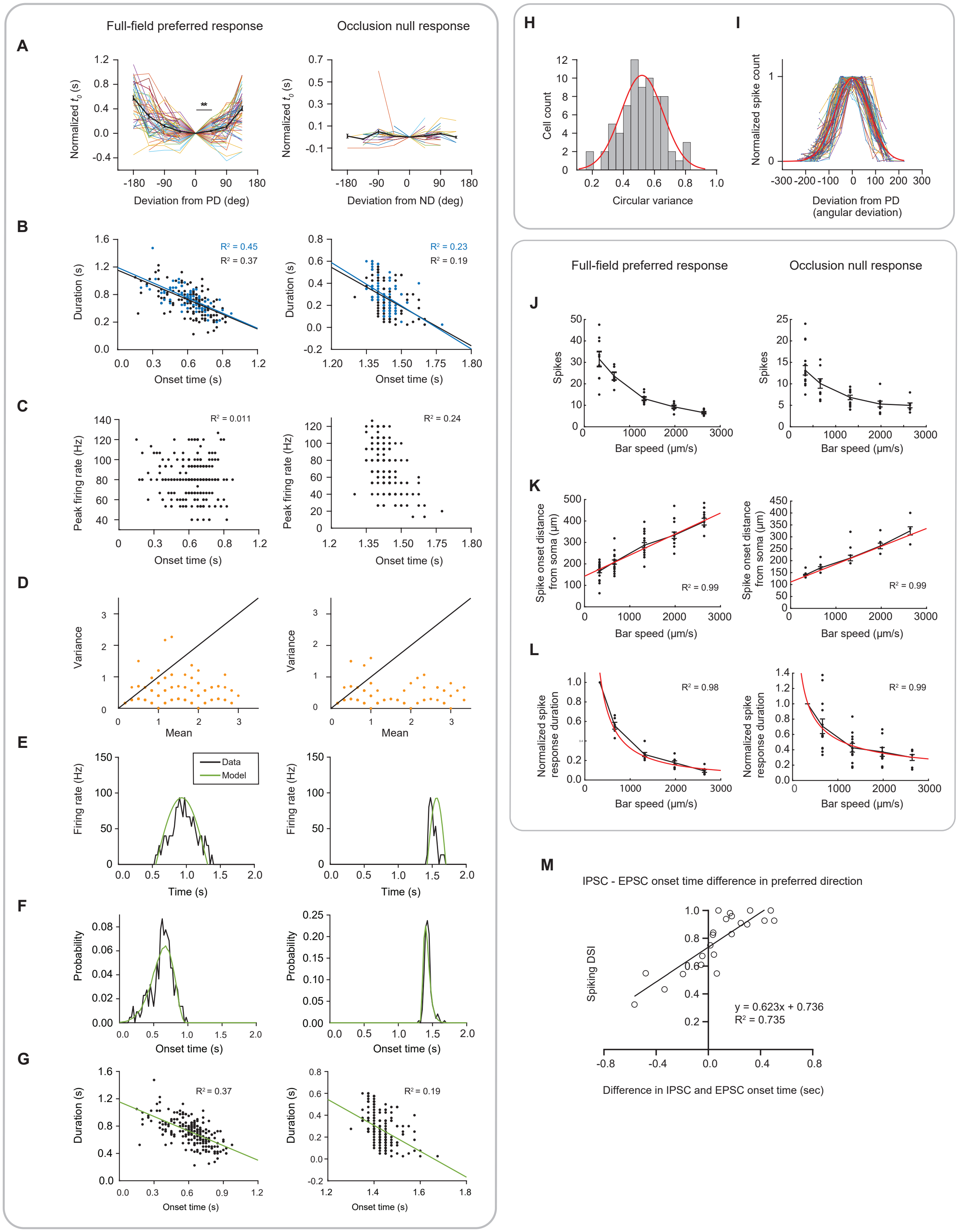


**Supplemental Figure 7. Model fits to experimental recordings.**
**(A)** Experimentally measured, normalized onset times, $t_0$, as a function of motion direction, with the average shown in bold and black (left: full-field, 73 cells; right: occlusion, 69 cells). Onset times were aligned to the time when the bar moved in the cell's preferred direction (full-field

preferred response 0 vs 45: **p = 0.004). **(B)** Linear correlation between spike response duration and spike response onset time. Data points are trials in which the bar moved in the preferred (left, full-field response) or null (right, occlusion response) direction (blue) or ± 45 degrees around the preferred or null direction (black). Left: full-field (black slope = -0.71; blue slope = -0.73). Right: occlusion (black slope = -1.18; blue slope = -1.31). **(C)** Scatter plots of peak firing rate versus onset time. Left: full-field; right: occlusion. **(D)** Scatter plots of spike count mean and variance (left: full-field, 10 cells; right: occlusion, 9 cells). The unity line is plotted. The spiking is significantly sub-Poisson. **(E)** Example half sine wave fits (green) to PSTHs obtained from experimental data (left: full-field, 73 cells; right: occlusion, 69 cells). **(F)** Fits to probability distributions of spike response onset times. **(G)** Linear correlation between spike response duration and spike response onset time. **(H)** Distribution of tuning curve widths fit to Gaussian. **(I)** Motion direction tuning curves of all cells normalized by their peaks and widths and their average (bold, red). Summary statistics are mean ± SEM. **(J)** Spiking response across speeds to the full-field preferred response (right, 10 cells) and the occlusion null response (left, 17 cells). **(K)** Same as **J** but for the spike onset distance from soma. **(L)** Same as J but for the normalized spike response duration. **(M)** The onset time difference between the IPSC and EPSC to a full-field bar in the preferred direction versus direction selectivity.
